# Immunization with synthetic SARS-CoV-2 S glycoprotein virus-like particles protects Macaques from infection

**DOI:** 10.1101/2021.07.26.453755

**Authors:** Guidenn Sulbaran, Pauline Maisonnasse, Axelle Amen, Delphine Guilligay, Nathalie Dereuddre-Bosquet, Judith A. Burger, Meliawati Poniman, Marlyse Buisson, Sebastian Dergan Dylon, Thibaut Naninck, Julien Lemaître, Wesley Gros, Anne-Sophie Gallouët, Romain Marlin, Camille Bouillier, Vanessa Contreras, Francis Relouzat, Daphna Fenel, Michel Thepaut, Isabelle Bally, Nicole Thielens, Franck Fieschi, Guy Schoehn, Sylvie van der Werf, Marit J. van Gils, Rogier W. Sanders, Pascal Poignard, Roger Le Grand, Winfried Weissenhorn

## Abstract

The SARS-CoV-2 pandemic causes an ongoing global health crisis, which requires efficient and safe vaccination programs. Here, we present synthetic SARS-CoV2 S glycoprotein-coated liposomes that resemble in size and surface structure virus-like particles. Soluble S glycoprotein trimers were stabilized by formaldehyde cross-linking and coated onto lipid vesicles (S-VLP). Immunization of cynomolgus macaques with S-VLPs induced high antibody titers and TH1 CD4+ biased T cell responses. Although antibody responses were initially dominated by RBD specificity, the third immunization boosted non-RBD antibody titers. Antibodies showed potent neutralization against the vaccine strain and the Alpha variant after two immunizations and robust neutralization of Beta and Gamma strains. Challenge of animals with SARS-CoV-2 protected all vaccinated animals by sterilizing immunity. Thus, the S-VLP approach is an efficient and safe vaccine candidate based on a proven classical approach for further development and clinical testing.

## Introduction

SARS-CoV-2, a betacoronavirus closely related to SARS-CoV-1 is the etiological agent of coronavirus disease (COVID-19), which quickly developed into a worldwide pandemic (Coronaviridae Study Group of the International Committee on Taxonomy of, 2020; Zhou et al., 2020) causing more than 4 million deaths as of July 2021 (https://covid19.who.int/) and highlighting the urgent need for effective infection control and prevention.

Antiviral vaccines achieve protection by generating neutralizing antibodies. The main SARS-CoV-2 target for inducing neutralizing antibodies is the S glycoprotein composed of the S1 subunit that harbors the receptor-binding domain (RBD) and the S2 membrane fusion subunit that anchors the S trimer in the virus membrane (Dagotto et al., 2020). RBD binding to the cellular receptor Angiotensin-converting enzyme 2 (ACE2) induces virus uptake and subsequent S2-mediated fusion with endosomal membranes establishes infection (Letko et al., 2020; Tortorici and Veesler, 2019; Wrapp et al., 2020). S is synthesized as a trimeric precursor polyprotein that is proteolytically cleaved by furin and furin-like protease in the Golgi generating the non-covalently linked S1-S2 heterotrimer (Hoffmann et al., 2020). The structure of S reveals a compact heterotrimer composed of S1 NTD, RBD, RBM and two subdomains, S2, the transmembrane region and a cytoplasmic domain. The conformation of RBD is in a dynamic equilibrium between either all RBDs in a closed, receptor-inaccessible conformation or one or two RBDs in the “up”, receptor-accessible, conformation (Cai et al., 2020; Ke et al., 2020; Turonova et al., 2020; Walls et al., 2020b; Wrapp et al., 2020). Only S RBD in the ‘up’ position allows receptor binding (Lan et al., 2020; Yan et al., 2020), which triggers the S2 post fusion conformation in proteolytically cleaved S (Cai et al., 2020). S is also highly glycosylated, which affects infection (Thépaut et al., 2021) and access to neutralizing antibodies (Watanabe et al., 2020).

Antibodies targeting the S glycoprotein were identified upon SARS-CoV-2 seroconversion (Amanat et al., 2020), which mostly target RBD that is immunodominant (Piccoli et al., 2020; Premkumar et al., 2020). This led to the isolation of many neutralizing antibodies, which confirmed antibody-based vaccination strategies (Baum et al., 2020; Brouwer et al., 2020; Hansen et al., 2020; Kreer et al., 2020; Liu et al., 2020a; Pinto et al., 2020; Robbiani et al., 2020; Rogers et al., 2020; Seydoux et al., 2020a; Seydoux et al., 2020b; Wang et al., 2020a; Wec et al., 2020; Wu et al., 2020; Yuan et al., 2020; Zost et al., 2020). Many of these antibodies have been shown to provide *in vivo* protection against SARS-Cov-2 challenge in small animals and nonhuman primates (Alsoussi et al., 2020; Hassan et al., 2020; Tortorici et al., 2020; Wu et al., 2020; Zost et al., 2020) or are in clinical development (Weinreich et al., 2021).

The magnitude of antibody responses to S during natural infection varies greatly and correlates with disease severity and duration (Chen et al., 2020; Seow et al., 2020b). Basal responses are generally maintained for months (Isho et al., 2020b; Iyer et al., 2020; Rodda et al., 2020) or decline within weeks after infection (Seow et al., 2020a), which is even faster in asymptomatic individuals (Long et al., 2020). Thus, any vaccine-based approach aims to induce long-lasting immunity.

A number of animal models have been developed to study SARS CoV-2 infection including the macaque model, which demonstrated induction of innate, cellular and humoral responses upon infection (Grifoni et al., 2020; Maisonnasse et al., 2020; McMahan et al., 2020; Munster et al., 2020; Rockx et al., 2020) conferring partial protection against reinfection (Deng et al., 2020; Marlin et al., 2021). Consequently, many early vaccine candidates provided protection in the macaque model including the currently licensed vaccines based on S-specific mRNA delivery (Corbett et al., 2020; Vogel et al., 2021) (BNT162b2, Pfizer/BioNTech; mRNA-1273, Moderna), adenovirus vectors (Mercado et al., 2020; van Doremalen et al., 2020) (ChAdOx1 nCoV-19, Oxford/AstraZeneca; *Ad26*.COV2.S, Johnson & Johnson) and inactivated SARS-CoV-2 (Gao et al., 2020; Wang et al., 2020b) (PiCoVacc/CoronaVac, Sinovac). Numerous other approaches have been evaluated as well (Klasse et al., 2021).

Employing the classical subunit approach, S subunit vaccine candidates have generated different levels of neutralizing antibody responses in preclinical testing (Liang et al., 2021; Liu et al., 2020b; Mandolesi et al., 2021; Tan et al., 2021a; Zang et al., 2020). Employing self-assembly strategies of S or RBD further increased immune responses (Walls et al., 2020a; Zhang et al., 2020a) and protected against infection (Arunachalam et al., 2021; Brouwer et al., 2021; Guebre-Xabier et al., 2020).

Antigens can be also presented via liposomes, which provide a high controllable degree of multivalency, stability and prolonged circulating half-life *in vivo* (Alving et al., 2016; Nisini et al., 2018). Notably, liposomes coated with viral glycoproteins such as HIV-1 Env induced more efficient immune responses than immunization with single glycoprotein trimers (Bale et al., 2017; Dubrovskaya et al., 2019; Ingale et al., 2016; Martinez-Murillo et al., 2017). This is in line with a more efficient B cell activation and generation of germinal centers (GC) by multivalent presentation of Env trimers versus soluble trimers (Ingale et al., 2016).

Here we developed synthetic virus-like particles employing liposomes that are decorated with S glycoprotein trimers that have been treated by formaldehyde cross linking, which in turn stabilized S in the native conformation over a long time-period. Serum antibody recognition of cross-linked versus non-cross-linked S did not show significant binding differences. A small group of cynomolgus macaques were immunized with S-VLPs, which produced high S antibody titers TH1 CD4+ T cell responses. Potent neutralization of wild-type SARS-CoV2 (WT) and of Alpha pseudovirus variants is observed after two immunizations, while Beta and Gamma pseudovirus variants are neutralized at reduced potency. Challenge of the animals with SARS-CoV-2 demonstrated that S-VLP immunization protected the animals from infection revealing no detection of genomic RNA (gRNA) upon infection in nasal and tracheal swabs, nor in broncho-alveolar lavages (BAL), thus providing sterilizing immunity. This indicates that S-VLPs are potential candidates for further clinical development of a safe protein-based SARS CoV-2 vaccine.

## Results

### S-VLP formation and characterization

The S glycoprotein construct ‘2P’ (Wrapp et al., 2020) was expressed in mammalian cells and purified by Ni^2+^-affinity and size exclusion chromatography (SEC) (**Figure S1A**), with yields up to 10 mg/liter using Expi293F cells. This produced native trimers as determined by negative staining electron microscopy and 2-D class averaging of the single particles (**Figure 1A**). Since S revealed low thermostability (Tm = 42°C) as reported previously (Wrapp et al., 2020), it was chemically cross-linked with 4% formaldehyde (FA) producing a higher molecular weight species as determined by SDS-PAGE (**Figure 1A**). FA cross-linking preserved the native structure over longer time periods (**Figures 1B, C and D**) by increasing the thermostability to a Tm of 65 °C. FA cross-linked S (FA-S) was incubated with liposomes (Phosphatidylcholine 60%, Cholesterol 36%, DGS-NTA 4%), and efficiently captured via its C-terminal His-tag. Free, unbound S was removed from the S proteoliposomes by sucrose gradient centrifugation (**Figure S1B**) and decoration of the liposomes with FA-S (S-VLP) was confirmed by negative staining electron microscopy (**Figure 1E**).

**Figure 1:**
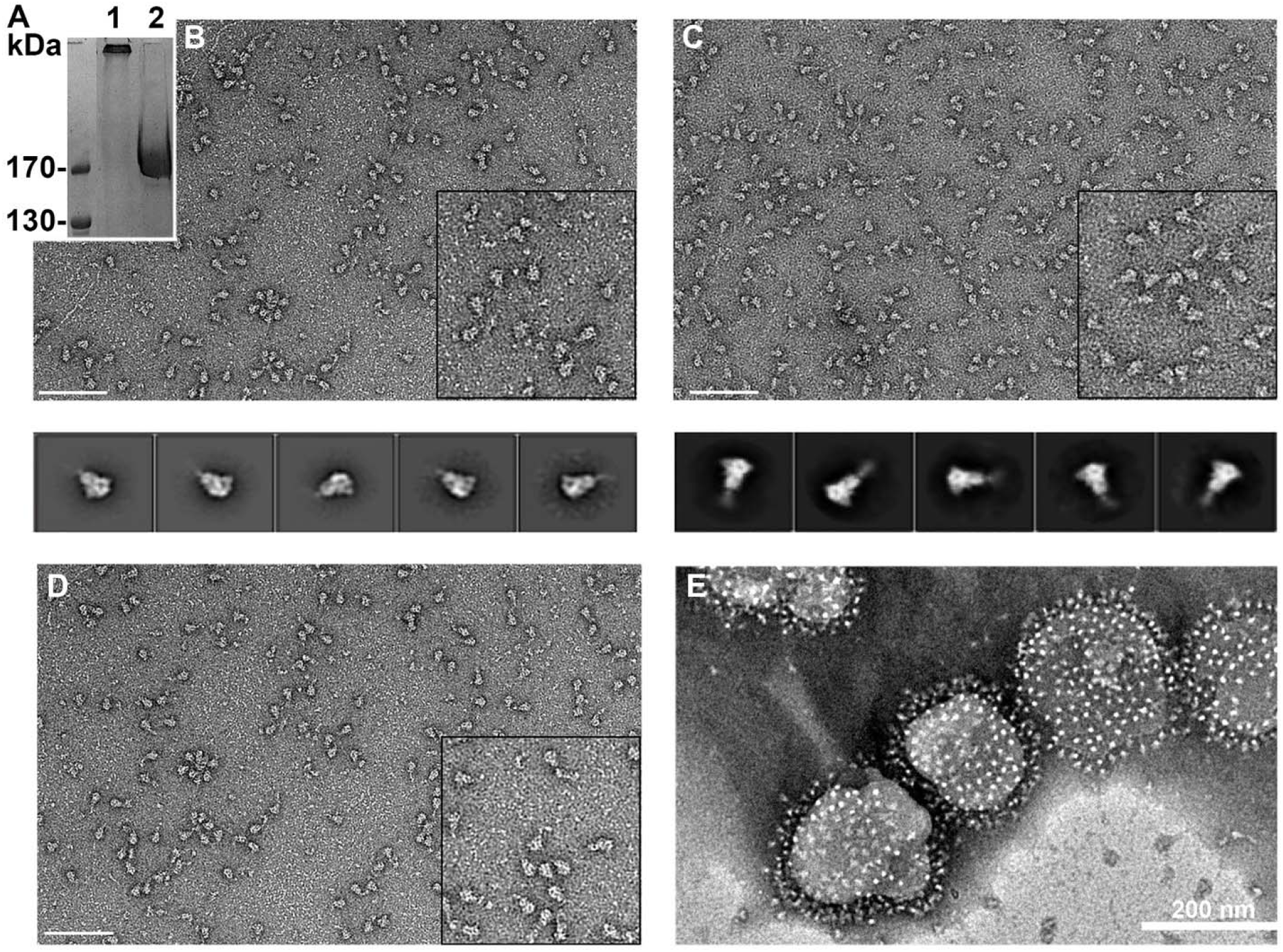
Production of SARS CoV-2 S glycoprotein. **(A)** SDS-PAGE of purified SARS-CoV-2 S (lane 2) and S chemically cross-linked with 4% formaldehyde (FA) (Lane 1). **(B)** Negative staining electron microscopy of the S glycoprotein before and **(C)** after FA cross linking. 2-D class averages of the five most populated classes are shown below the panels, which indicate native closed trimers. **(D)** FA-cross-linked S glycoprotein analyzed after storage (2 weeks) at 4°C. The lower panels in B and C show representative 2-D class averages. Scale bars are 200 nm. **(E)** FA-cross-linked S glycoprotein was incubated with liposomes containing 4% DGS-NTA lipids, purified by sucrose gradient density centrifugation and analyzed by negative staining electron microscopy revealing regular decoration of the liposomes with the S glycoprotein. Scale bar, 200 nm.

### S-VLP immunization induces potent neutralizing antibody responses in cynomolgus macaques

S-VLPs were produced for a small vaccination study of cynomolgus macaques to evaluate safety, immunogenicity and elicitation of neutralizing antibodies. Four cynomolgus macaques were immunized with 50 μg S-VLPs adjuvanted with monophospholipid A (MPLA) liposomes by the intramuscular route at weeks 0, 4, 8 and 19 (**Figure 2A**). Sera of the immunized macaques were analysed for binding to native S glycoprotein (S), FA cross-linked S glycoprotein (FA-S) and RBD in two weeks intervals. This revealed median S ED50 titers of 100 at week 4, 3000 at week 8 and 25 000 at week 12 (**Figure 2B**). Slight reductions in titers were detected against FA-S (**Figure 2C**). Titers against RBD alone were also high with median ED50s of 80 at week 4, 2000 at week 8 and 5200 at week 12 (**Figure 2D**). This suggests that the first and second immunization induced significant RBD titers, while the third immunization boosted non-RBD antibodies since the week 12 S-specific titers were > 5 times higher than the RBD titers (**Figure 2C**). A fourth immunization did not further boost antibody generation and titers at week 22 were lower or comparable to week 12 titers (**Figure 2B, C, D**). We conclude that S-VLP immunization induces primarily RBD-specific antibodies after the first and second immunization, while the third immunization increases the generation of non-RBD antibodies significantly.

**Figure 2:**
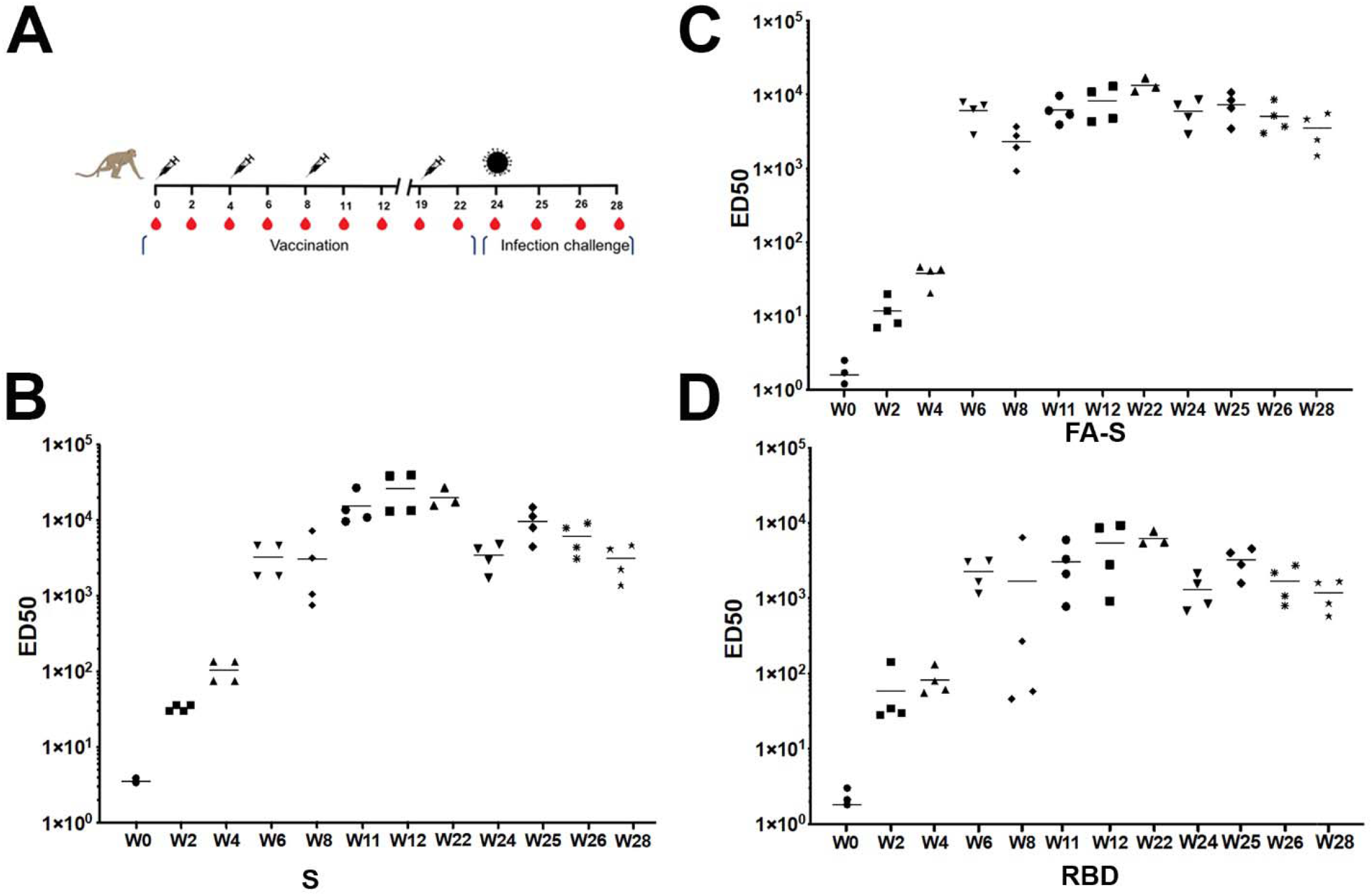
Antibody responses induced by S-VLP vaccination of cynomolgus macaques. **(A)** Scheme of vaccination, challenge and sampling. Syringes indicate the time points of vaccination, red drops the time of serum collection and the virus particle the time point of challenge. Symbols of identifying individual macaques are used in all figures. **(B)** ELISA of SARS-CoV-2 S-protein-specific IgG determined during the study at weeks 0, 2, 4, 6, 8, 11, 12, 19, 22, 24, 25, 28. Median values calculated for the 4 animals are indicated. **(C)** ELISA of SARS-CoV-2 FA-S-protein-specific IgG determined during the study. **(D)** ELISA of SARS-CoV-2 S RBD-specific IgG determined during the study.

Serum neutralization titers using WT pseudovirus were significant in all four animals. At week 2 after the first immunization, a median ID50 titer of 480 was observed, which dropped close to baseline at week 4, but was significantly increased at week 6, two weeks after the second immunization demonstrating a median ID50 of 9,000. The ID50s then decreased to a median of 6,000 at week 8 and increased to a median of 18,500 at week 11, three weeks after the third immunization. At week 19, neutralization potency decreased but was still high with a median of 5,200, indicating that three immunizations induced robust neutralization titers. The fourth immunization boosted neutralization titers to a median ID50 of 20,000, the same level as after the third immunization (**Figure 3A**).

**Figure 3:**
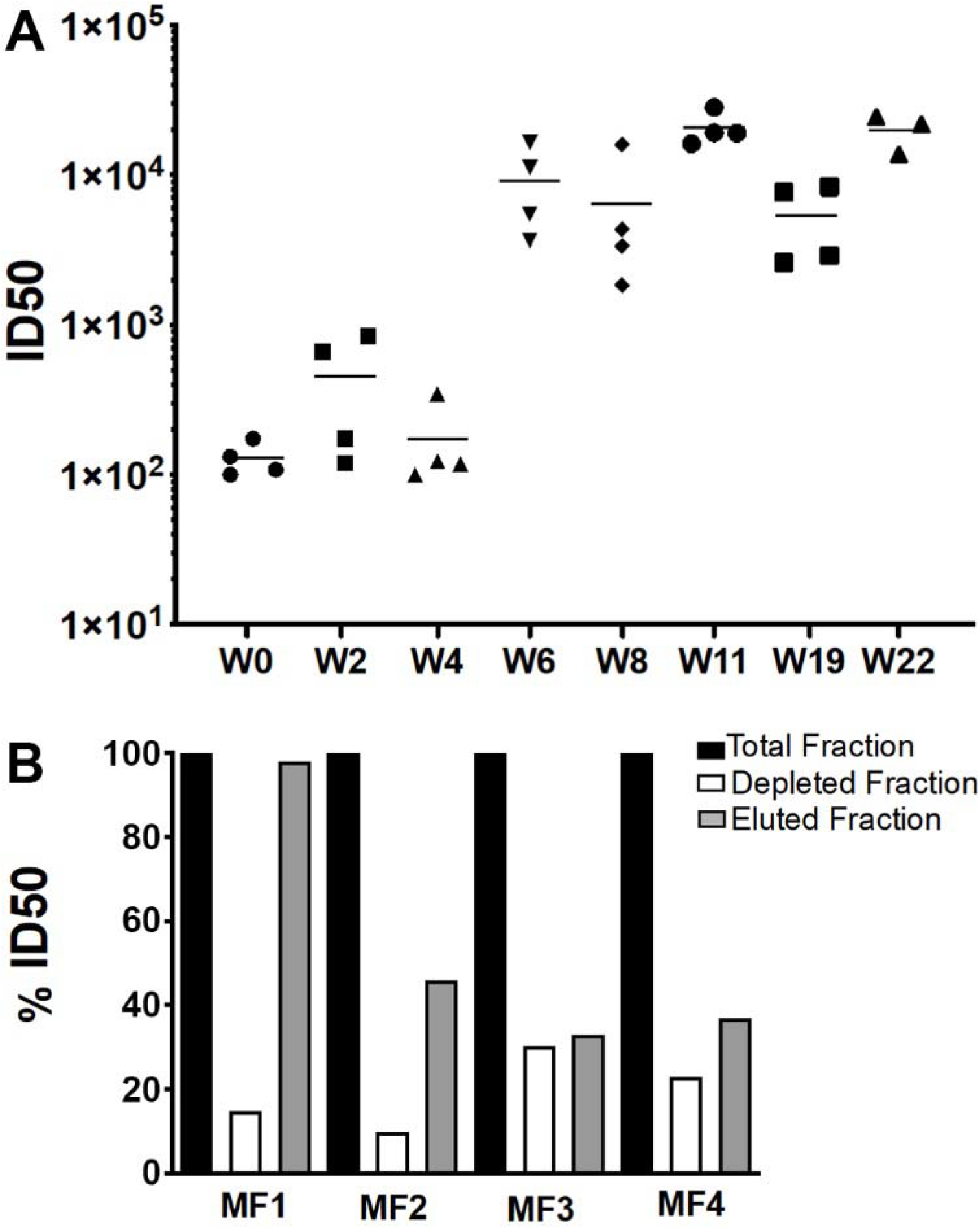
Serum neutralization of SARS-CoV-2 pseudovirus upon S-VLP vaccination. **(A)** The evolution of SARS-CoV-2 neutralizing Ab titers is shown for sera collected at weeks 0, 2, 4, 6, 8, 11, 12, 19. Bars indicate median titers of the four animals. B) Serum from week 11 was depleted of RBD-specific Abs by affinity chromatography and neutralization activity of the complete serum of each animal was set to 100 % and compared to the RBD-depleted sera and the RBD-specific sera.

Since antibody titers indicated the induction of high levels of RBD-specific antibodies, we depleted the serum at week 11 by anti-RBD affinity chromatography resulting in no detectable RBD antibodies by ELISA. RBD-specific Ab-depleted serum showed 10 to 30% neutralization compared to the complete serum, indicating non-RBD specific neutralization. While RBD-specific Ab neutralization dominated in one animal, the other three revealed 30 to 48% RBD-specific Ab neutralization activity (**Figure 3B**), suggesting nAb synergy to achieve the high neutralization titers (**Figure 3A**).

### S-VLP immunization protects cynomolgus macaques from SARS-CoV-2 infection

In order to determine the extent of S-VLP vaccination induced protection, vaccinated and non-vaccinated animals (n=4) were infected with the primary SARS-CoV-2 isolate (BetaCoV/France/IDF/0372/2020) with a total dose of 1 × 10^5^ plaque forming units (pfu). Infection was induced by combining intranasal (0.25 mL into each nostril) and intratracheal (4.5 mL) routes at week 24, 5 weeks after the last immunization. Viral load in the control animal group peaked in the trachea at 3 days post-exposure (dpe) with a median value of 6.0 log_10_ copies/ml and in the nasopharynx at day 6 pe with a median copy number of 6,6 log_10_ copies/ml (**Figure 4A**). Viral loads decreased subsequently and no virus was detected on day 10 pe in the trachea, while some animals showed viral detection up to day 14 pe in the nasopharyngeal swabs (**Figure 4A**). In the bronchoalveolar lavage (BAL), three CTRL animal out of four showed detectable viral loads at day 3 pe, and two of them remained detectable at day 7 pe with mean value of 5.4 and 3.6 log_10_ copies/mL respectively. Rectal fluids tested positive in one animal, which also had the highest tracheal and nasopharyngeal viral loads (**Figure S2**). Viral subgenomic RNA (sgRNA), which is believed to estimate the number of infected and productive cells collected with the swabs or during the lavage, showed peak copy numbers between day 3/4 and 6 pe in the tracheal and nasopharyngeal fluids, respectively (**Figure 4B**). In the BALs, the two animals presenting high genomic viral loads also showed detectable sgRNA at days 3 and 7 pe, with medians of 5,1 and 3.1 log_10_ copies/mL respectively (**Figure 4B**).

**Figure 4:**
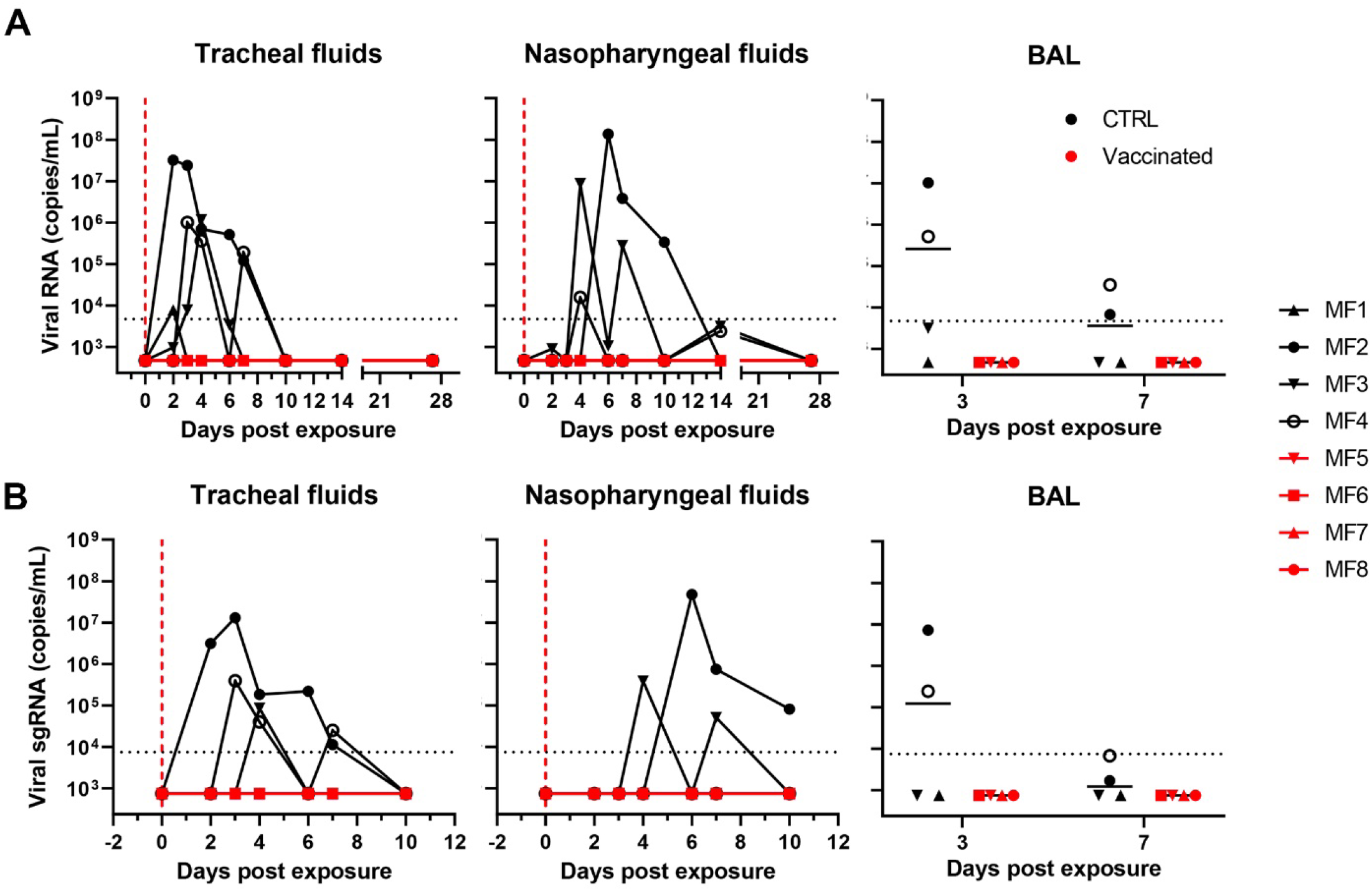
S-VLP immunization protects cynomolgus macaques from SARS-CoV-2 infection. (A) RNA viral loads in tracheal swabs (left) and nasopharyngeal swabs (middle) of control and vaccinated macaques after challenge. Viral loads in control and vaccinated macaques after challenge in BAL are shown (right). Bars indicate median viral loads. Vertical red dotted lines indicate the day of challenge. Horizontal dotted lines indicate the limit of quantification. (B) sgRNA viral loads in tracheal swabs (left), nasopharyngeal swabs (middle), and BAL (right) of control and vaccinated macaques after challenge. Bars indicate mean viral loads. Dotted line indicates the limit of quantification. The symbols identifying individual animals are indicated.

Neither gRNA nor sgRNA was detected at any point in the vaccinated group (**Figures 4A, B**), indicating sterilizing immunity induced by vaccination, both in the upper and lower respiratory tract. In line with this observation, no increase in Ab titers and neutralization was observed in the vaccinated animals. Median ID50 antibody titers against S, FA-S and RBD decreased from 8000 to 3000 from the day of infection to 3 weeks pe (**Figure 5A, B, C**), while the control group started to show a slight increase in IgG Abs against RBD after week 1 and a clear detection of S and FA-S IgG from week 2 on (**Figure 5A, B, C**). Consistent with the IgG Ab responses, neutralization activity continued to decrease from the day of challenge at week 24 to week 28 from a median ID50 of 10,000 (week 24) to 7,000 (week 28) (**Figure 5E**). This demonstrated that challenge of vaccinated animals did not boost their immune system. In contrast non-vaccinated animals showed significant neutralization at week 2 pe with a median ID50 of 900 (week 26) followed by the observation of a decline of neutralization titers up to week 32, week 8 pe (**Figure 5D**).

**Figure 5:**
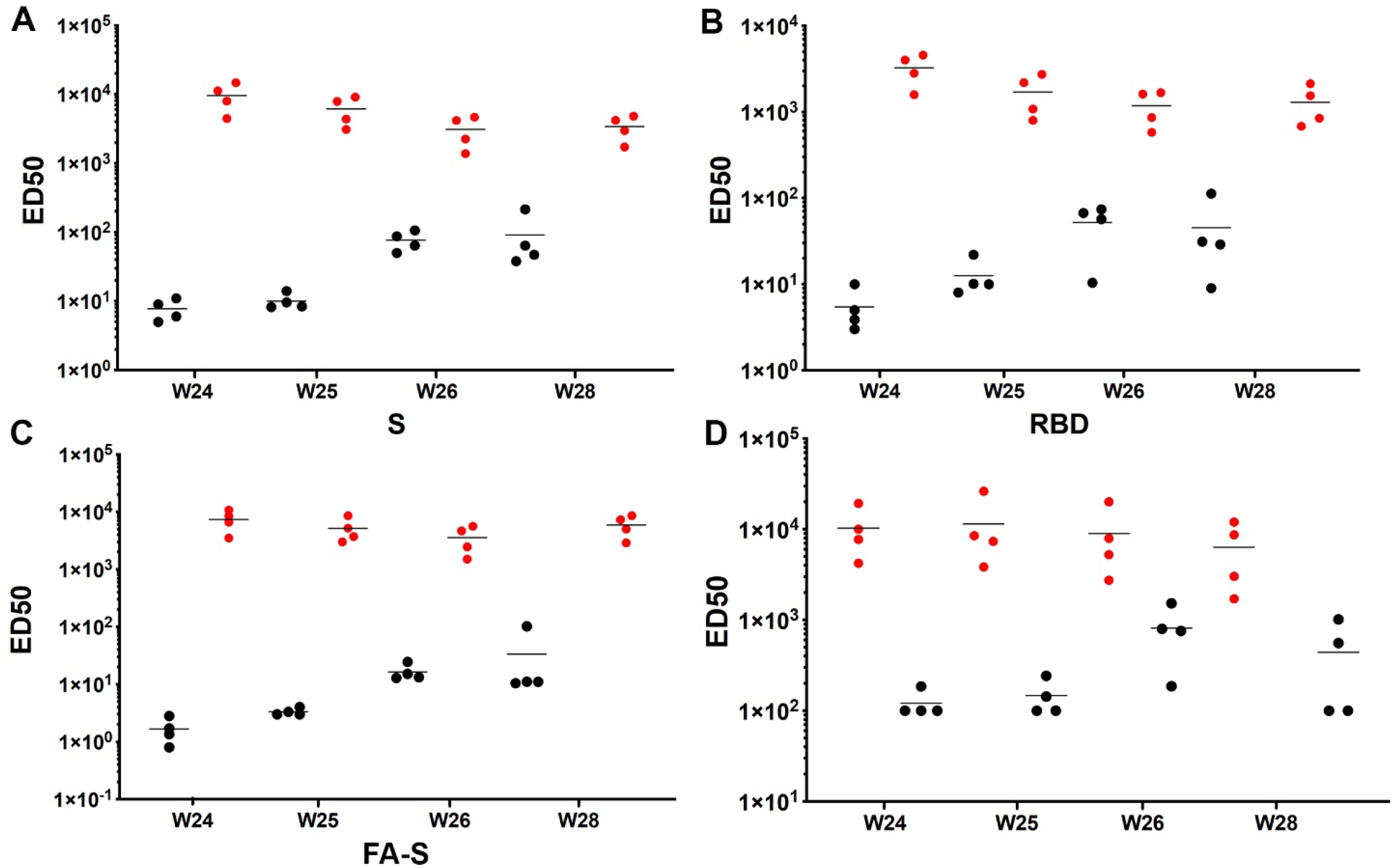
Serum antibody titers and neutralization of vaccinated and control group cynomolgus macaques after SARS CoV-2 challenge. Antibody IgG titers were determined by ELISA at weeks 24 (challenge), 25, 26 and 28 against **(A)** SARS-CoV-2 S, **(B)** SARS-CoV-2 FA-S and **(C)** SARS-CoV-2 S RBD. Vaccinated animals are shown with red symbols and control animals with black symbols. **(D)** SARS CoV-2 pseudovirus neutralization titers at week 24 (challenge) and 1, 2 and 4 weeks post exposure (weeks 25, 26, 28). The Bars show the median titers.

Similar to previous observations (Maisonnasse et al., 2020; Brouwer et al., 2021), during the first 14 dpe, all control animals showed mild pulmonary lesions characterized by nonextended ground-glass opacities (GGOs) detected by chest computed tomography (CT) (**Figure S3A**). Vaccinated animals showed no significant impact of challenge on CT scores. The only animal showing a lesion score >10 was in the CTRL group. Whereas all control animals experienced monocytoses between days 2 and 8 pe, probably corresponding to a response to infection, monocytes counts remained stable after challenge for the vaccinated monkeys (**Figure S3B**), in agreement with the absence of detectable anamnestic response.

Before exposure, Th1 type CD4^+^ T-cell responses were observed in all vaccinated macaques following *ex vivo* stimulation of PBMCs with S-peptide pools (**Figure 6 and S6**). None had detectable anti-S CD8^+^ T cells (**Figure S5**). No significant difference was observed at day 14 pe, also in agreement with the absence of an anamnestic response. In contrast, the anti-S Th1 CD4+ response increased post exposure for most of the control animals (**Figure S6 and S7**).

**Figure 6:**
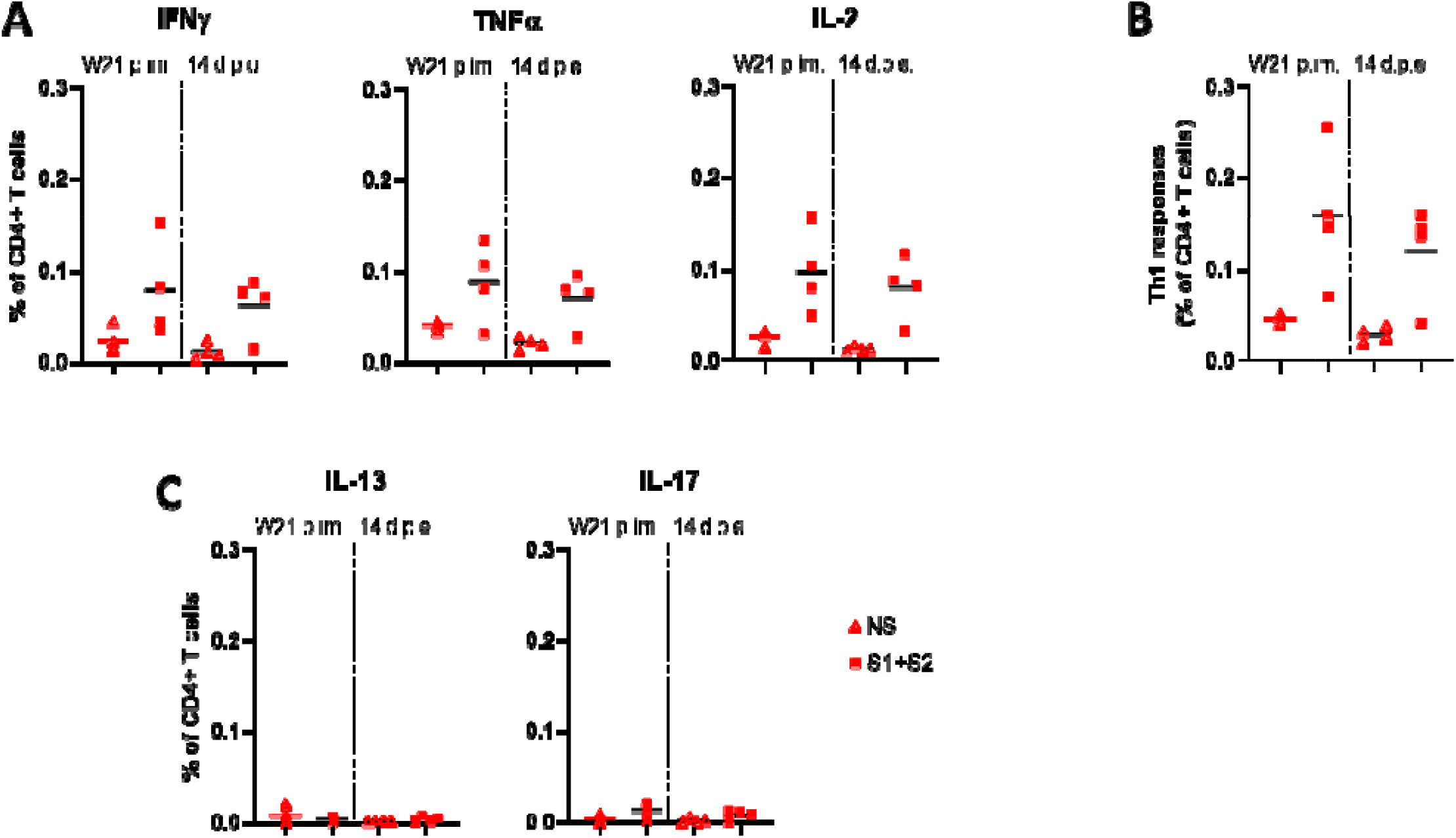
Antigen-specific CD4 T-cell responses in S-VLP immunized cynomolgus macaques. Frequency of (A) IFNγ+, TFNα+ and IL-2+, (B), Th1 (IFN γ+/−, IL-2+/−, TNFα+), (C) IL-13+and IL-17+ antigen-specific CD4+ T cells (CD154+) in the total CD4+ T cell population, respectively, for each immunized macaque (n = 4) at week (W)21 post-immunization (p.im.) (i.e. two weeks after the 4^th^ immunization, pre-exposure) and 14 days post-exposure (d.p.e.). PBMCs were stimulated overnight with medium (open symbols) or SARS-CoV-2 S overlapping peptide pools (filled symbols). Time points in each experimental group were compared using the Wilcoxon signed rank test.

We conclude that S-VLP vaccination can produce sterilizing immunity indicating that the vaccination scheme is efficient to interrupt the chain of transmission.

### S-VLP vaccination generates robust neutralization of SARS-CoV-2 variants

Serum neutralization was further tested against variants B.1.1.7 (Alpha, UK), B.1.351 (Beta, SA) and P.1 (Gamma, BR). Comparing the sera of the vaccinated and the non-vaccinated group at weeks 24 and 28 showed high neutralization titers for all three variants with median ID50s ranging from 10.000 to 20.000, comparable to WT pseudovirus neutralization (**Figure S4**). However, since the background of serum neutralization of the non-vaccinated challenge group was relatively high (median ID50s ranging from 400 of 1100), we repeated the neutralization with purified IgG from serum samples of the vaccinated group from week 8 (after 2 immunizations), week 12 (3 immunizations) and weeks 24 and 28 (4 immunizations). This showed median ID50s of 2500 for WT and Alpha on week 8, which indicated that IgG purification reduced the potency by a factor of 2,6 compared to complete serum neutralization (**Figure 3A**). Lower ID50 medians of 150 and 450 were observed against Beta and Gamma at week 8, respectively. Neutralization potency was increased after the third immunization (week 12) with median ID50s of 2000 (WT), 6000 (Alpha), 500 (Beta) and 1000 (Gamma). Neutralization titers did not increase after the fourth immunization at week 24 and started to decrease at week 28 (**Figure 7**). We conclude that three immunizations provide robust protection against the variants although neutralization titers maybe already within the protective range after two immunizations for the three variants tested.

**Figure 7:**
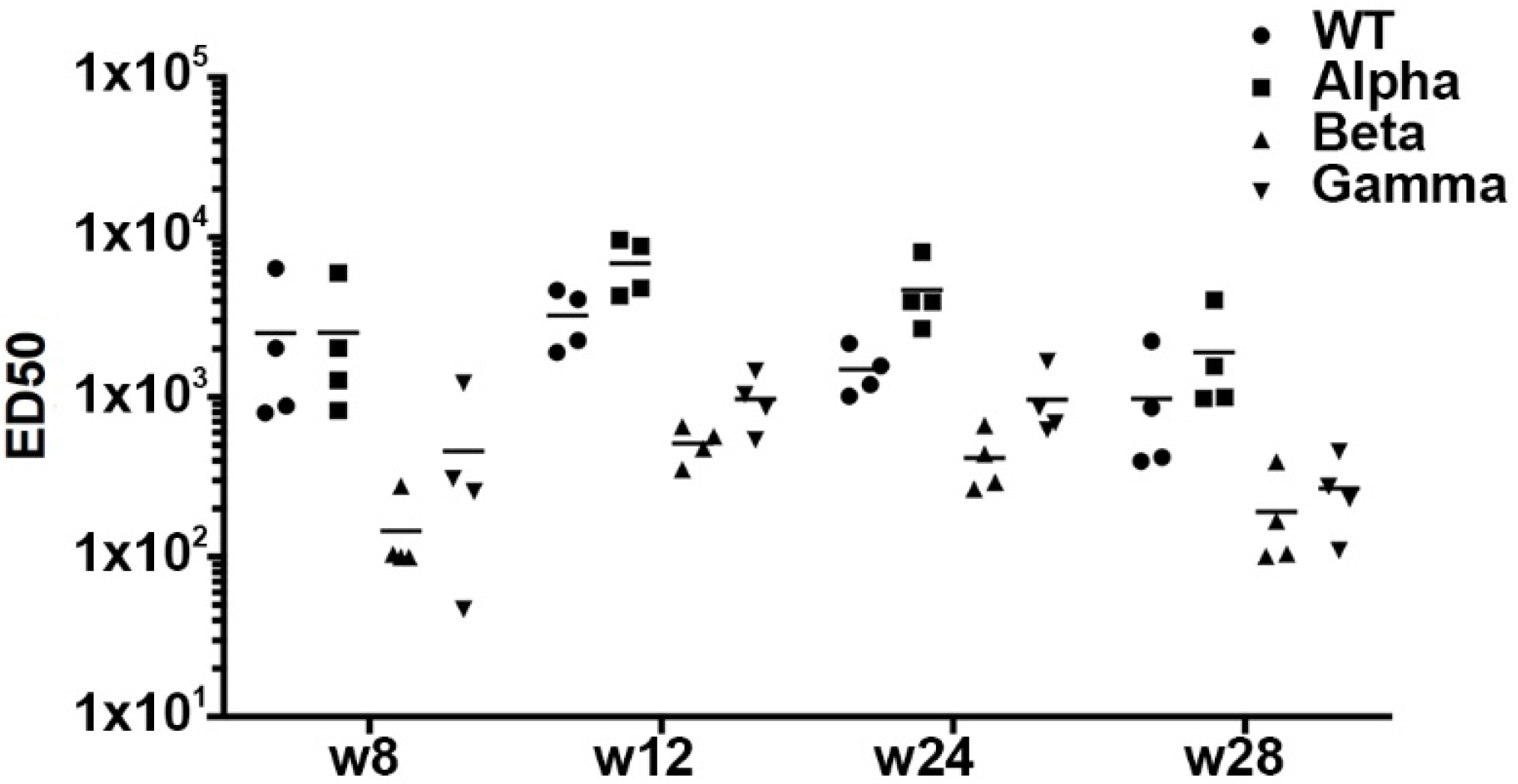
S-VLP vaccination induces robust neutralization of SARS CoV-2 variants. B.1.1.7 (Alpha, UK), B.1.351 (Beta, SA) and P.1 (Gamma, BR) pseudovirus neutralization titers were compared to the Wuhan vaccine strain. Titers were determined using total IgG purified from sera at weeks 8 (2 immunizations), 12 (3 immunizations), 24 and 28 (4 immunizations).

## Discussion

Many vaccines are under development, in preclinical and clinical testing (Klasse et al., 2021) and eight have been approved by regulatory agencies over the world. Here we developed a two-component system employing SARS-CoV-2 S glycoprotein coupled to liposomes. Since the stability of the wild type SARS-CoV-2 S glycoprotein is low due to its tendency to spontaneously switch into its post fusion conformation (Cai et al., 2020), SARS-CoV-2 S was stabilized by two proline mutations that enhanced stability (Wrapp et al., 2020). However, this S ‘2P’ version still showed limited stability over time as reported (Wrapp et al., 2020), which may be due to cold sensitivity (Edwards et al., 2021). We overcame the problem of stability by using formaldehyde cross-linking that increased the thermostability to 65°C, preserving the native S conformation over extended storage time periods. Notably, formaldehyde cross-linking is widely used in vaccine formulations (Eldred et al., 2006). S stability has been since improved by engineering six proline mutations (S ‘6P’) which increased the thermostability to 50°C (Hsieh et al., 2020) and by disulfide-bond engineering (Xiong et al., 2020). Furthermore, ligand binding renders S more stable (Rosa et al., 2021; Toelzer et al., 2020).

Many previous studies have shown that immunogen multimerization strategies are highly beneficial for B cell activation including the use of synthetic liposomes (Alving et al., 2016; Nisini et al., 2018) such as HIV-1 Env-decorated liposome vaccination strategies (Dubrovskaya et al., 2019). We linked SARS-CoV-2 S to liposomes producing synthetic virus-like particles with controlled diameters. These S-VLPs show similar immunogenic properties as a number of reported self-assembling particles of SARS-CoV-2 RBD and S (Brouwer et al., 2021; Cohen et al., 2021; Guebre-Xabier et al., 2020; Tan et al., 2021b; Walls et al., 2020a; Zhang et al., 2020a). Our S-VLPs induce robust and potent neutralizing responses in cynomolgus macaques, which completely protected the animals from infection by sterilizing immunity. Notably, no signs of virus replication could be detected in the upper and lower respiratory tracts consistent with the absence of clinical signs of infection such as lymphopenia and lung damage characteristics for Covid-19 disease. The important correlate of protection against SARS-CoV-2 is provided by neutralizing antibodies (Addetia et al., 2020; McMahan et al., 2020; Yu et al., 2020). The S-VLP approach induces high titers already after two immunizations, with a median ID50 of 6000 four weeks after the second immunization, which is substantially higher than neutralizing Ab responses reported for vaccines tested in NHP studies, including licensed ones. Adenovirus-based vaccines (AstraZeneca ChAdOx1; Janssen AD26COV2SPP°(Mercado et al., 2020; van Doremalen et al., 2020), inactivated virus vaccines (Sinovac PiCoVacc; Sinopharm/BIBP BBIP-CorV)(Gao et al., 2020; Wang et al., 2020b), DNA vaccine (Yu et al., 2020) and a mRNA vaccine (Pfizer/BionTech BNT162b2)(Vogel et al., 2021) showed 10-20 times lower titers compared to the S-VLPs. Moderna mRNA-1273 (Moderna)(Corbett et al., 2020), S trimers (Clover Biopharmaceutical) (Liang et al., 2021) and NVX-CoV2373 (Novavax)(Guebre-Xabier et al., 2020) induced similar or higher titer. Median ID50 titers increased by a factor of ~4 after the third immunization, but did not amplify after the fourth immunization. The T cell response in the vaccinated group was biased towards TH1 CD4+ T cells consistent with licensed or other experimental vaccines (Corbett et al., 2020; Ewer et al., 2021; Ganneru et al., 2021; Keech et al., 2020; Sahin et al., 2020).

Serum neutralization was already significant after the first immunization, but increased by a factor of ~20 after the second immunization and by a factor of 3 after the third immunization indicating that two immunizations with S-VLPs may suffice to confer protection. BnAb titers decline within 11 weeks after the third immunization to the levels of week 8 (prior to the third immunization) and increase to the median ID50 level attained after the third immunization. ID 50 neutralization values decline by a factor of ~4 between week 22 and week 28 after the fourth immunization consistent with general Ab decline over time.

Vaccination prevented lymphopenia and lung damage in animals infected with SARS-CoV-2 at a dose comparable (Corbett et al., 2020; Guebre-Xabier et al., 2020; Mercado et al., 2020; van Doremalen et al., 2020; Yu et al., 2020) or lower (Brouwer et al., 2021) than in previous studies. Protection was sterilizing since no replication could be detected in the upper and lower respiratory tract suggesting that vaccination with S-VLPs will prevent virus shedding and transmission. Sterilizing immunity likely correlates with mucosal antibody responses that protects the upper respiratory tract from infection (Isho et al., 2020a; Randad et al., 2020). However, we failed to detect significant IgA or IgG in nasopharyngeal fluids, which may be due to the low sensitivity of the ELISA tests used.

Most of the antibodies (up to 90%) generated by vaccination are directed against RBD, which is immunodominant (Piccoli et al., 2020). RBD antibodies can be grouped into three classes (Barnes et al., 2020a; Barnes et al., 2020b) and seem to be easily induced by immunization as many of them are generated by few cycles of affinity maturation indicating that extensive germinal center reactions are not required (Kreye et al., 2020). Consistent with these findings we show that RBD-specific antibodies are predominant after the first and second immunization revealing similar S-specific and RBD-specific titers. However, after the third immunization median S-specific ED50s are 3 times higher than RBD-specific ED50s four weeks after the third immunization. This trend is continued after the fourth immunization which revealed a 3.5 times higher median ID50 for S than for RBD five weeks post immunization. This, thus demonstrates that more than two immunizations allow to expand the reactive B cell repertoire that target non-RBD S epitopes.

Current variants carry the B.1 D614G mutation and have been reported to be more infectious (Cai et al., 2021; Gobeil et al., 2021; Korber et al., 2020; Ozono et al., 2021; Yuan et al., 2021; Zhang et al., 2021; Zhang et al., 2020b). Although the D614G mutation alone was reported to increase neutralization susceptibility (Weissman et al., 2021), further mutations present in Beta (B1.351 SA) and Gamma (P1, BR) reduce neutralization potencies of natural and vaccine-induced sera (Dejnirattisai et al., 2021; Edara et al., 2021; Garcia-Beltran et al., 2021; Geers et al., 2021; Hoffmann et al., 2021; Kuzmina et al., 2021; Rees-Spear et al., 2021; Zhou et al., 2021), while Alpha (B.1.1.7, UK) neutralization seems to be less affected (Supasa et al., 2021). Reduction in neutralization potency of polyclonal plasma Abs is mainly affected by mutations within the three main epitopes in the RBD and especially the E484K mutation present in Beta and Gamma was reported to reduce neutralization by a factor of 10 (Greaney et al., 2021). Here we show that S-VLP vaccination produces robust neutralization of Alpha, Beta and Gamma although the Median ID50s of Beta and Gamma neutralization are reduced 20- and 5-fold after the second immunization compared to WT and Alpha. The third immunization boosted neutralization of Beta and Gamma, albeit with 6-fold (Beta) and 3-fold (Gamma) reductions in potency compared to WT, which is slightly more potent than the median ID50 of vaccinated and hospitalized patient cohorts using the same assay setup (Caniels et al., 2021) without taking into account that serum IgG purification reduced the ID50 of the WT ~threefold.

In summary, S-VLP vaccination represents an efficient strategy that protects macaques from high dose challenge. Although the animals have been challenged only after the fourth immunization, which did not boost Ab titers or neutralization titers, our neutralization data suggests that the animals might have been protected after two immunizations. Furthermore, we provide evidence that the third immunization boosts non-RBD antibodies which is likely important to protect against different variants. This also suggests that future vaccination strategies should probably boost non-RBD antibodies to compensate for the loss of neutralization targeting RBD. Notably, SARS-CoV-2 memory B cells are present over a long time period (Gaebler et al., 2021; Sokal et al., 2021), which is in line with different boosting strategies. Finally, although other regions within S, notably NTD are targets for mutation within new variants, S2 or other epitopes may be less prone to mutations due to conformational constraints.

## Methods

### Protein expression and purification

The SARS-CoV-2 S gene encoding residues 1-1208 with proline substitutions at residues 986 and 987 (“2P”), a “GSAS” substitution at the furin cleavage site (residues 682-685) a C-terminal T4 fibritin trimerization motif, an HRV3C protease cleavage site, a TwinStrepTag and Hexa-His-tag (McLellan et al., 2020) was transiently expressed in FreeStyle293F cells (Thermo Fisher scientific) using polyethylenimine (PEI) 1 μg/μl for transfection. Supernatants were harvested five days post-transfection, centrifuged for 30 min at 5000 rpm and filtered using 0.20 μm filters (ClearLine®). SARS-CoV-2 S protein was purified from the supernatant by Ni^2+^-Sepharose chromatography (Excel purification resin, Cytiva) in buffer A (50 mM HEPES pH 7.4, 200 mM NaCl) and eluted in buffer B (50 mM HEPES pH 7.4, 200 mM NaCl, 500 mM imidazole). Eluted SARS-CoV-2 S containing fractions were concentrated using Amicon Ultra (cut-off: 30 KDa) (Millipore) and further purified by size-exclusion chromatography (SEC) on a Superose 6 column (GE Healthcare) in buffer A or in PBS.

For RBD expression, the following reagent was produced under HHSN272201400008C and obtained through BEI Resources, NIAID, NIH: Vector pCAGGS containing the SARS-Related Coronavirus 2, Wuhan-Hu-1 Spike Glycoprotein Receptor Binding Domain (RBD), NR-52309. The SARS-CoV-2 S RBD domain (residues 319 to 541) was expressed in EXPI293 cells by transient transfection according to the manufacturer’s protocol (Thermo Fisher Scientific). Supernatants were harvested five days after transfection and cleared by centrifugation. The supernatant was passed through a 0.45 μm filter and RBD was purified using Ni^2+^-chromatography (HisTrap HP column, GE Healthcare) in buffer C (20 mM Tris pH 7.5 and 150 mM NaCl buffer) followed by a washing step with buffer D (20 mM Tris pH 7.5 and 150 mM NaCl buffer, 75 mM imidazole) and elution with buffer E (20 mM Tris pH 7.5 and 150 mM NaCl buffer, 500 mM imidazole). Eluted RBD was further purified by SEC on a Superdex 75 column (GE Healthcare) in buffer C. Protein concentrations were determined using an absorption coefficient (A1%,1cm) at 280 nm of 10.4 and 13.06 for S protein and RBD, respectively, using the ProtParam available at https://web.expasy.org/.

### SARS-CoV-2 S crosslinking

S protein at 1mg/ml in PBS was cross-linked with 4% formaldehyde (FA) (Sigma) overnight at room temperature. The reaction was stopped with 1 M Tris HCl pH 7.4 adjusting the sample concentration to 7.5 mM Tris/HCl pH 7.4. FA was removed by PBS buffer exchange using 30 KDa cut-off concentrators (Amicon). FA crosslinking was confirmed by separating SARS-CoV-2 FA-S on a 10% SDS-PAGE under reducing conditions.

### S protein coupling to liposomes

Liposomes for conjugating S protein were prepared as described previously (Scianimanico et al., 2000) with modifications. Briefly, liposomes were composed of 60% of L-α-phosphatidylcholine, 4% His tag-conjugating lipid, DGS-NTA-(Ni^2+^) and 36% cholesterol (Avanti Polar Lipids). Lipid components were dissolved in chloroform, mixed and placed for two hours in a desiccator under vacuum at room temperature to obtain a lipid film. The film was hydrated in filtered (0.22 μm) PBS and liposomes were prepared by extrusion using membrane filters with a pore size of 0.1 μm (Whatman Nuclepore Track-Etch membranes). The integrity and size of the liposomes was analyzed by negative staining-EM. For protein coupling, the liposomes were incubated overnight with FA-S or S protein in a 3:1 ratio (w/w). Free FA-S protein was separated from the FA-S-proteoliposomes (S-VLPs) by sucrose gradient (5-40%) centrifugation in a SW55 rotor at 40,000 rpm for 2 h. The amount of protein conjugated to the liposomes was determined by Bradford assay and SDS-PAGE densitometry analysis comparing S-VLP bands with standard S protein concentrations.

### S protein thermostability

Thermal denaturation of SARS-CoV-2 S, native or FA-cross-linked was analyzed by differential scanning fluorimetry coupled to back scattering using a Prometheus NT.48 instrument (Nanotemper Technologies, Munich, DE). Protein samples were first extensively dialyzed against PBS pH 7.4, and the protein concentration was adjusted to 0.3 mg/ml. 10 μl of sample were loaded into the capillary and intrinsic fluorescence was measured at a ramp rate of 1°C/min with an excitation power of 30 %. Protein unfolding was monitored by the changes in fluorescence emission at 350 and 330 nm. The thermal unfolding midpoint (Tm) of the proteins was determined using the Prometheus NT software.

### Negative stain electron microscopy

Protein samples were visualized by negative-stain electron microscopy (EM) using 3-4 μl aliquots containing 0.1-0.2 mg/ml of protein. Samples were applied for 10 s onto a mica carbon film and transferred to 400-mesh Cu grids that had been glow discharged at 20 mA for 30 s and then negatively stained with 2% (wt/vol) Uranyl Acetate (UAc) for 30 s. Data were collected on a FEI Tecnai T12 LaB6-EM operating at 120 kV accelerating voltage at 23k magnification (pixel size of 2.8 Å) using a Gatan Orius 1000 CCD Camera. Two-dimensional (2D) class averaging was performed with the software Relion (Scheres, 2012) using on average 30–40 micrographs per sample. The 5 best obtained classes were calculated from around 6000 particles each.

### Ethics and biosafety statement

Cynomolgus macaques (Macaca fascicularis) originating from Mauritian AAALAC certified breeding centers were used in this study. All animals were housed in IDMIT infrastructure facilities (CEA, Fontenay-aux-roses), under BSL-2 and BSL-3 containment when necessary (Animal facility authorization #D92-032-02, Préfecture des Hauts de Seine, France) and in compliance with European Directive 2010/63/EU, the French regulations and the Standards for Human Care and Use of Laboratory Animals, of the Office for Laboratory Animal Welfare (OLAW, assurance number #A5826-01, US). The protocols were approved by the institutional ethical committee “Comité d’Ethique en Expérimentation Animale du Commissariat à l’Energie Atomique et aux Energies Alternatives” (CEtEA #44) under statement number A20-011. The study was authorized by the “Research, Innovation and Education Ministry” under registration number APAFIS#24434-2020030216532863.All information on the ethics committee is available at https://cache.media.enseignementsup-recherche.gouv.fr/file/utilisation_des_animaux_fins_scienifiques/22/1/comiteethiqueea17_juin2013_257221.pdf.

### Viruses and cells

For the macaques studies, SARS-CoV-2 virus (hCoV-19/France/ lDF0372/2020 strain) was isolated by the National Reference Center for Respiratory Viruses (Institut Pasteur, Paris, France) as previously described (Lescure et al., 2020) and produced by two passages on Vero E6 cells in DMEM (Dulbecco’s Modified Eagles Medium) without FBS, supplemented with 1% P/S (penicillin at 10,000 U ml^−1^ and streptomycin at 10,000 μg ml^−1^) and 1 μg ml^−1^ TPCK-trypsin at 37 °C in a humidified CO_2_ incubator and titrated on Vero E6 cells. Whole genome sequencing was performed as described (Lescure et al., 2020) with no modifications observed compared with the initial specimen and sequences were deposited after assembly on the GISAID EpiCoV platform under accession number ID EPI_ISL_410720.

### Animals and study design

Cynomolgus macaques were randomly assigned in two experimental groups. The vaccinated group (n = 4) received 50 ug of SARSCoV-2 S-VLP adjuvanted with 500 mg of MPLA liposomes (Polymun Scientific, Klosterneuburg, Austria) diluted in PBS at weeks 0, 4, 8 and 19, while control animals (n = 4) received no vaccination. Vaccinated animals were sampled in blood at weeks 0, 2, 4, 6, 8, 11, 12, 14, 19, 21 and 22. At week 24, all animals were exposed to a total dose of 10^5^ pfu of SARS-CoV-2 virus (hCoV-19/France/ lDF0372/2020 strain; GISAID EpiCoV platform under accession number EPI_ISL_410720) via the combination of intranasal and intra-tracheal routes (0,25 mL in each nostril and 4,5 mL in the trachea, i.e., a total of 5 mL; day 0), using atropine (0.04 mg/kg) for pre-medication and ketamine (5 mg/kg) with medetomidine (0.042 mg/kg) for anesthesia. Nasopharyngeal, tracheal and rectal swabs, were collected at days 2, 3, 4, 6, 7, 10, 14 and 27 days past exposure (dpe) while blood was taken at days 2, 4, 7, 10, 14 and 27 dpe. Bronchoalveolar lavages (BAL) were performed using 50 mL sterile saline on 3 and 7 dpe. Chest CT was performed at 3, 7, 10 and 14 dpe in anesthetized animals using tiletamine (4 mg kg^−1^) and zolazepam (4 mg kg^−1^). Blood cell counts, haemoglobin, and haematocrit, were determined from EDTA blood using a DHX800 analyzer (Beckman Coulter).

### Virus quantification in NHP samples

Upper respiratory (nasopharyngeal and tracheal) and rectal specimens were collected with swabs (Viral Transport Medium, CDC, DSR-052-01). Tracheal swabs were performed by insertion of the swab above the tip of the epiglottis into the upper trachea at approximately 1.5 cm of the epiglottis. All specimens were stored between 2°C and 8°C until analysis by RT-qPCR with a plasmid standard concentration range containing an RdRp gene fragment including the RdRp-IP4 RT-PCR target sequence. SARS-CoV-2 E gene subgenomic mRNA (sgRNA) levels were assessed by RT-qPCR using primers and probes previously described (Corman et al., 2020; Wolfel et al., 2020): leader-specific primer sgLeadSARSCoV2-F CGATCTCTTGTAGATCTGTTCTC, E-Sarbeco-R primer ATATTGCA GCAGTACGCACACA and E-Sarbeco probe HEX-ACACTAGCCATCCTTACTGCGCTTCG-BHQ1. The protocol describing the procedure for the detection of SARS-CoV-2 is available on the WHO website^58^.

### Chest CT and image analysis

Lung images were acquired using a computed tomography (CT) system (Vereos-Ingenuity, Philips) as previously described (Brouwer et al., 2021; Maisonnasse et al., 2020), and analyzed using INTELLISPACE PORTAL 8 software (Philips Healthcare). All images had the same window level of −300 and window width of 1,600. Lesions were defined as ground glass opacity, crazy-paving pattern, consolidation or pleural thickening as previously described (Pan et al., 2020; Shi et al., 2020). Lesions and scoring were assessed in each lung lobe blindly and independently by two persons and the final results were established by consensus. Overall CT scores include the lesion type (scored from 0 to 3) and lesion volume (scored from 0 to 4) summed for each lobe as previously described (Brouwer et al., 2021; Maisonnasse et al., 2020).

### ELISA

Serum antibody titers specific for soluble native S glycoprotein, FA-cross-linked S (FA-S) and for RBD were determined using an enzyme-linked immunosorbent assay (ELISA). Briefly, 96-well microtiter plates were coated with 1 μg of S, FA-S or RBD proteins at 4°C overnight in PBS and blocked with 3% BSA for 1 h at room temperature after 3 washes with 150 μl PBS Tween-20 0.05 %. Serum dilutions were added to each well for 2h at 37°C and plates were washed 5 times with PBS Tween. A horseradish peroxidase (HRP) conjugated goat anti-monkey H+L antibody (Invitrogen #PA1-84631) was then added and incubated for 1h before excess Ab was washed out and HRP substrate added. Absorbance was determined at 450 nm. Antibody titers are presented as ED50 using the GraphPad Prism software version 6.

### Pseudovirus neutralization assay

Pseudovirus was produced by co-transfecting the pCR3 SARS-CoV-2-SΔ19 expression plasmid (Wuhan Hu-1; GenBank: MN908947.3) with the pHIV-1NL43 ΔEnv-NanoLuc reporter virus plasmid in HEK293T cells (ATCC, CRL-11268) (Caniels et al., 2021). The pCR3 SARS-CoV-2-SΔ19 expression plasmid contained the following mutations compared to the WT for the variants of concern: deletion (Δ) of H69, V70 and Y144, N501Y, A570D, D614G, P681H, T716I, S982A and D1118H in B.1.1.7; L18F, D80A, D215G, L242H, R246I, K417N, E484K, N501Y, D614G and A701V in B.1.351; L18F, T20N, P26S, D138Y, R190S, K417T, E484K, N501Y, D614G, H655Y and T1027I in P.1 (Caniels et al., 2021).

HEK293T/ACE2 cells kindly provided by Dr. Paul Bieniasz (Schmidt et al., 2020) were seeded at a density of 20,000 cells/well in a 96-well plate coated with 50 μg/mL poly-L-lysine 1 day prior to the start of the neutralization assay. Heat-inactivated sera (1:100 dilution) were serial diluted in 3-fold steps in cell culture medium (DMEM (Gibco), supplemented with 10% FBS, penicillin (100 U/mL), streptomycin (100 μg/mL) and GlutaMax (Gibco)), mixed in a 1:1 ratio with pseudovirus and incubated for 1 h at 37°C. These mixtures were then added to the cells in a 1:1 ratio and incubated for 48 h at 37°C, followed by a PBS wash and lysis buffer added. The luciferase activity in cell lysates was measured using the Nano-Glo Luciferase Assay System (Promega) and GloMax system (Turner BioSystems). Relative luminescence units (RLU) were normalized to the positive control wells where cells were infected with pseudovirus in the absence of sera. The neutralization titers (ID_50_) were determined as the serum dilution at which infectivity was inhibited by 50%, respectively using a non-linear regression curve fit (GraphPad Prism software version 8.3).

### Antigen specific T cell assays using non-human primate cells

To analyze the SARS-CoV-2 protein-specific T cell, 15-mer peptides (n = 157 and n=158) overlapping by 11 amino acids (aa) and covering the SARS-CoV-2 Spike sequence (aa 1 to 1273) synthesized by JPT Peptide Technologies (Berlin, Germany) and used at a final concentration of 2 μg/mL.

T-cell responses were characterized by measurement of the frequency of PBMC expressing IL-2 (PerCP5.5, MQ1-17H12, BD), IL-17a (Alexa700, N49-653, BD), IFN-γ (V450, B27, BD), TNF-α (BV605, Mab11, BioLegend), IL-13 (BV711, JES10-5A2, BD), CD137 (APC, 4B4, BD) and CD154 (FITC, TRAP1, BD) upon stimulation with the two peptide pools. CD3 (APC-Cy7, SP34-2, BD), CD4 (BV510, L200, BD) and CD8 (PE-Vio770, BW135/80, Miltenyi Biotec) antibodies was used as lineage markers. One million of PBMC were cultured in complete medium (RPMI1640 Glutamax+, Gibco; supplemented with 10 % FBS), supplemented with co-stimulatory antibodies (FastImmune CD28/CD49d, Becton Dickinson). Then cells were stimulated with S sequence overlapping peptide pools at a final concentration of 2 μg/mL Brefeldin A was added to each well at a final concentration of 10μg/mL and the plate was incubated at 37°C, 5% CO2 during 18 h. Next, cells were washed, stained with a viability dye (LIVE/DEAD fixable Blue dead cell stain kit, ThermoFisher), and then fixed and permeabilized with the BD Cytofix/Cytoperm reagent. Permeabilized cell samples were stored at −80 °C before the staining procedure. Antibody staining was performed in a single step following permeabilization. After 30 min of incubation at 4°C, in the dark, cells were washed in BD Perm/Wash buffer then acquired on the LSRII cytometer (Beckton Dickinson). Analyses were performed with the FlowJo v.10 software. Data are presented as the sum of each peptide pool and the non-stimulated (NS) condition was multiplied by two.

### Statistical analysis

Statistical analysis of NHP gRNA and sgRNA were carried out using Mann-Whitney unpaired t-test in GraphPad Prism software (v8.3.0).

## Acknowledgments

This work was supported by the the European Union’s Horizon 2020 research and innovation program under grant agreement No. 681032, H2020 EHVA (W.W.) and the ANR, RA-Covid-19. WW acknowledges access to the platforms of the Grenoble Instruct-ERIC center (IBS and ISBG; UMS 3518 CNRS-CEA-UGA-EMBL) within the Grenoble Partnership for Structural Biology (PSB), with support from FRISBI (ANR-10-INBS-05-02) and GRAL, a project of the University Grenoble Alpes graduate school (Ecoles Universitaires de Recherche) CBH-EUR-GS (ANR-17-EURE-0003). The IBS acknowledges integration into the Interdisciplinary Research Institute of Grenoble (IRIG, CEA) and financial support from CEA, CNRS and UGA. The Infectious Disease Models and Innovative Therapies (IDMIT) research infrastructure is supported by the Programme Investissements d’Avenir, managed by the National Research Agency (ANR) under reference ANR-11-INBS-0008. The Fondation Bettencourt Schueller and the Region Ile-de-France contributed to the implementation of IDMIT’s facilities and imaging technologies. The non-human primate study received financial support from REACTing, the Fondation pour la Recherche Médicale (AM-CoV-Path), and the European Infrastructure TRANSVAC2 (730964). We acknowledge support from CoVIC supported by the Bill and Melinda Gates Foundation. The virus stock was obtained through the EVAg platform (https://www.european-virus-archive.com/), funded by H2020 (653316). The funders had no role in study design, data collection, data analysis, data interpretation, or data reporting.

We thank J. McLellan for providing the S expression vector, B. Delache, E. Burban, J. Demilly, N. Dhooge, S. Langlois, P. Le Calvez, Q. Sconosciuti, V. Magneron, M. Rimlinger, A. Berriche, J.H. Qiu, M. Potier, J. M. Robert and C. Dodan for help with animal studies, and R. Ho Tsong Fang for his supervision; L. Bossevot, M. Leonec, L. Moenne-Loccoz, and J. Morin for the qRT-PCR and preparation of reagents; M. Gomez-Pacheco and J. van Wassenhove for cellular assays; N. Kahlaoui, B. Fert and C. Mayet for help with the CT scans, and C. Chapon for her supervision; M. Barendji, J. Dinh, and E. Guyon for the non-human primate sample processing; S. Keyser for the transports organization; F. Ducancel and Y. Gorin for their help with the logistics and safety management; and I. Mangeot for her help with resource management. We thank Antoine Nougairede for sharing the plasmid used for the sgRNA assay standardization. Finally, we thank Dietmar Katinger and Philipp Mundsperger (Polymun) for providing MPLA liposome adjuvants. Animal images in Figure 2 were created with BioRender.com.

---

**Figure S1:**
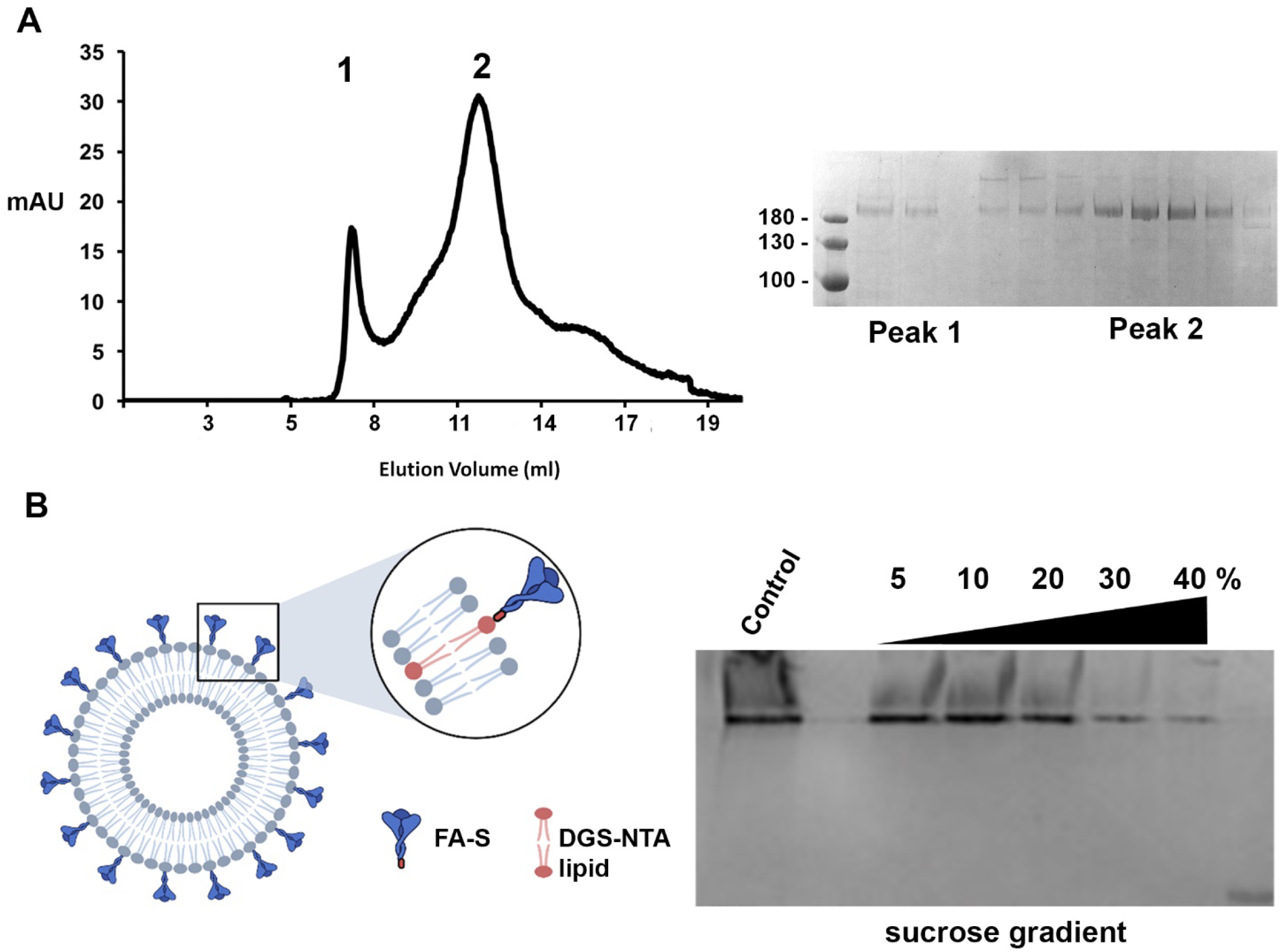
Expression and characterization of the SARS Cov-2 S glycoprotein. **(A)** SDS-PAGE and size exclusion chromatography (SEC) of purified S protein. **(B)** Right panel, schema of the liposome FA-S glycoprotein coupling; left panel, SDS-PAGE analysis of the S-VLP purification by sucrose gradient density centrifugation. S was incubated with liposomes containing DGS NTA lipids that capture FA-S via its C-terminal His-tag. Free FA-S was removed by sucrose gradient centrifugation.

**Figure S2:**
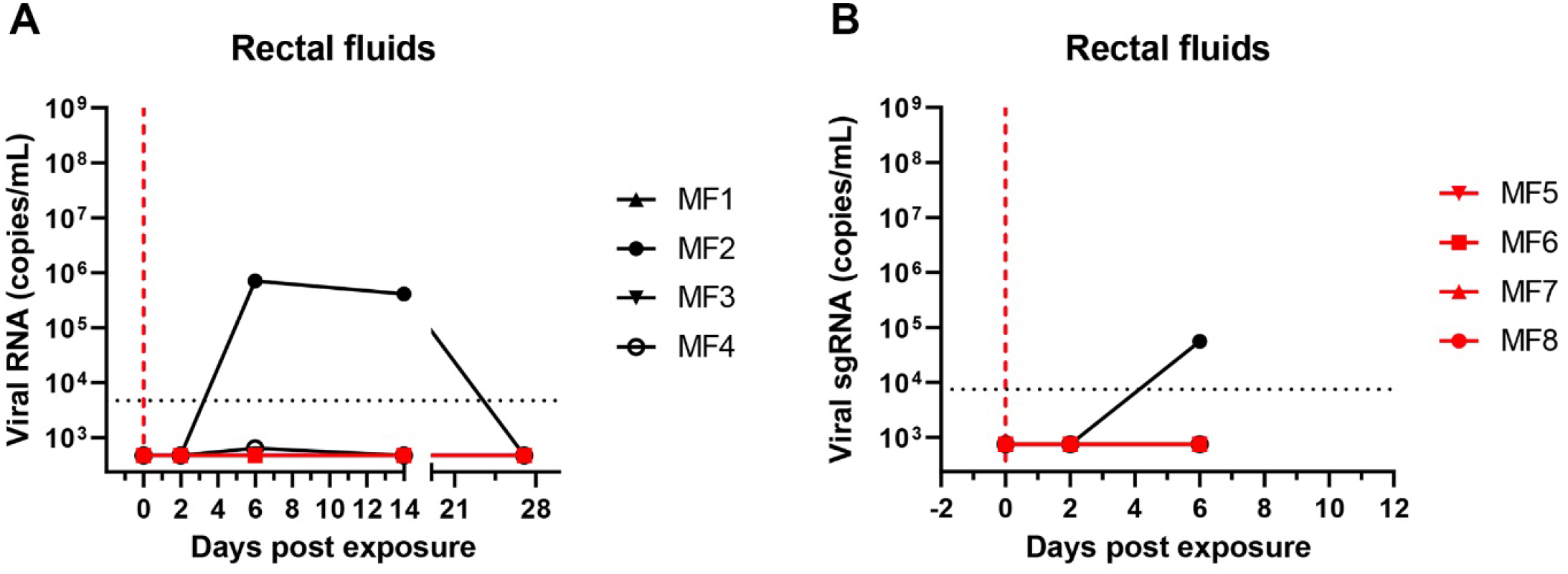
Viral RNA detection upon SARS CoV-2 challenge. (A) RNA viral loads detected in rectal fluids upon challenge of vaccinated animals and control animals. (B) Detection of viral sgRNA in rectal fluids. Data for individual animals (MF1-4, vaccinated group and MF5-8, control group) has been plotted.

**Figure S3:**
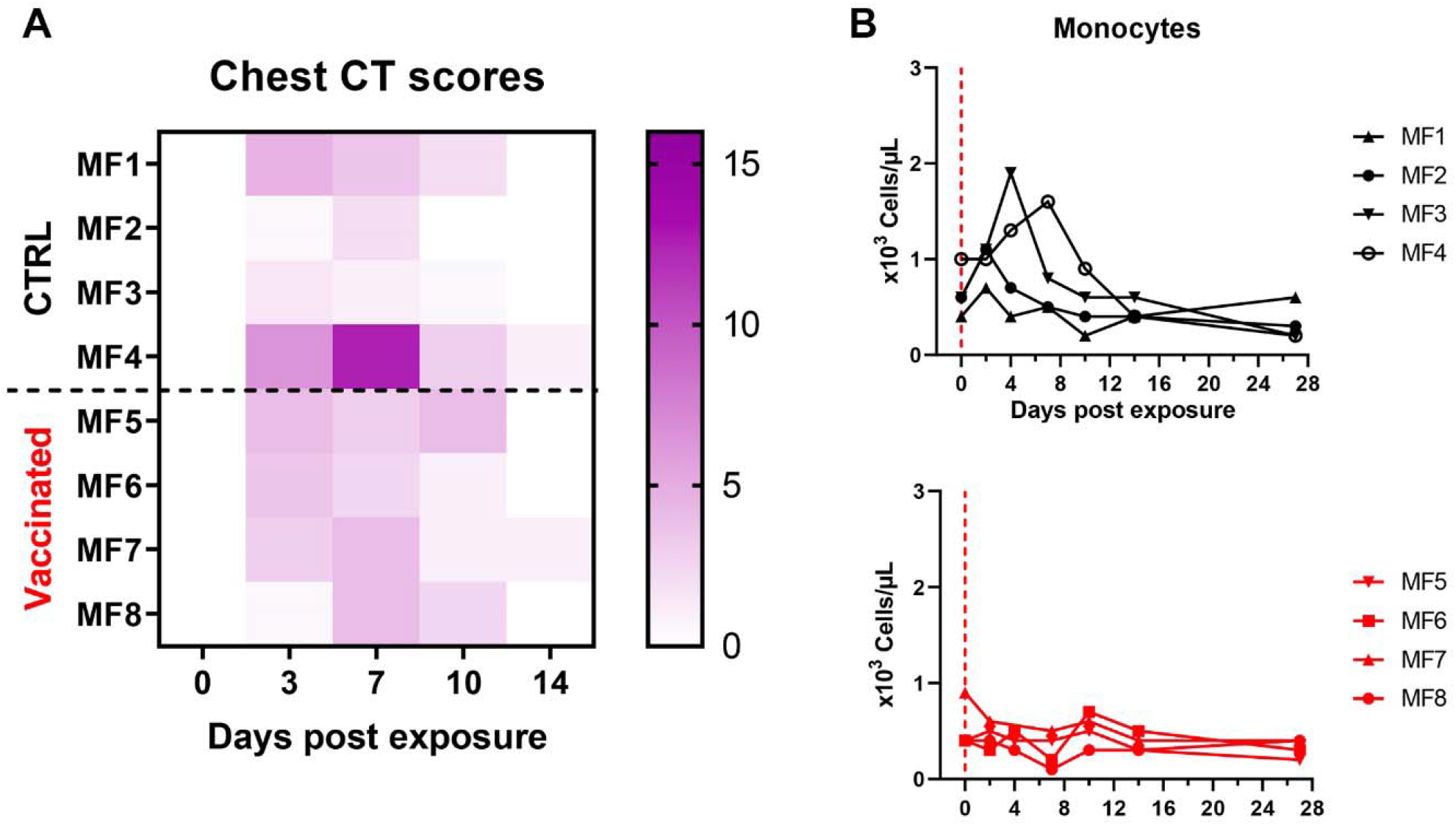
Clinical manifestations upon infection. (A) Lung CT scores of control and vaccinated macaques over the course of 14 days post exposure. The CT score is based on the lesion type (scored from 0 to 3) and lesion volume (scored from 0 to 4), which have been summed for each lobe. (B) Monocyte counts in the blood of control and vaccinated macaques after challenge.

**Figure S4:**
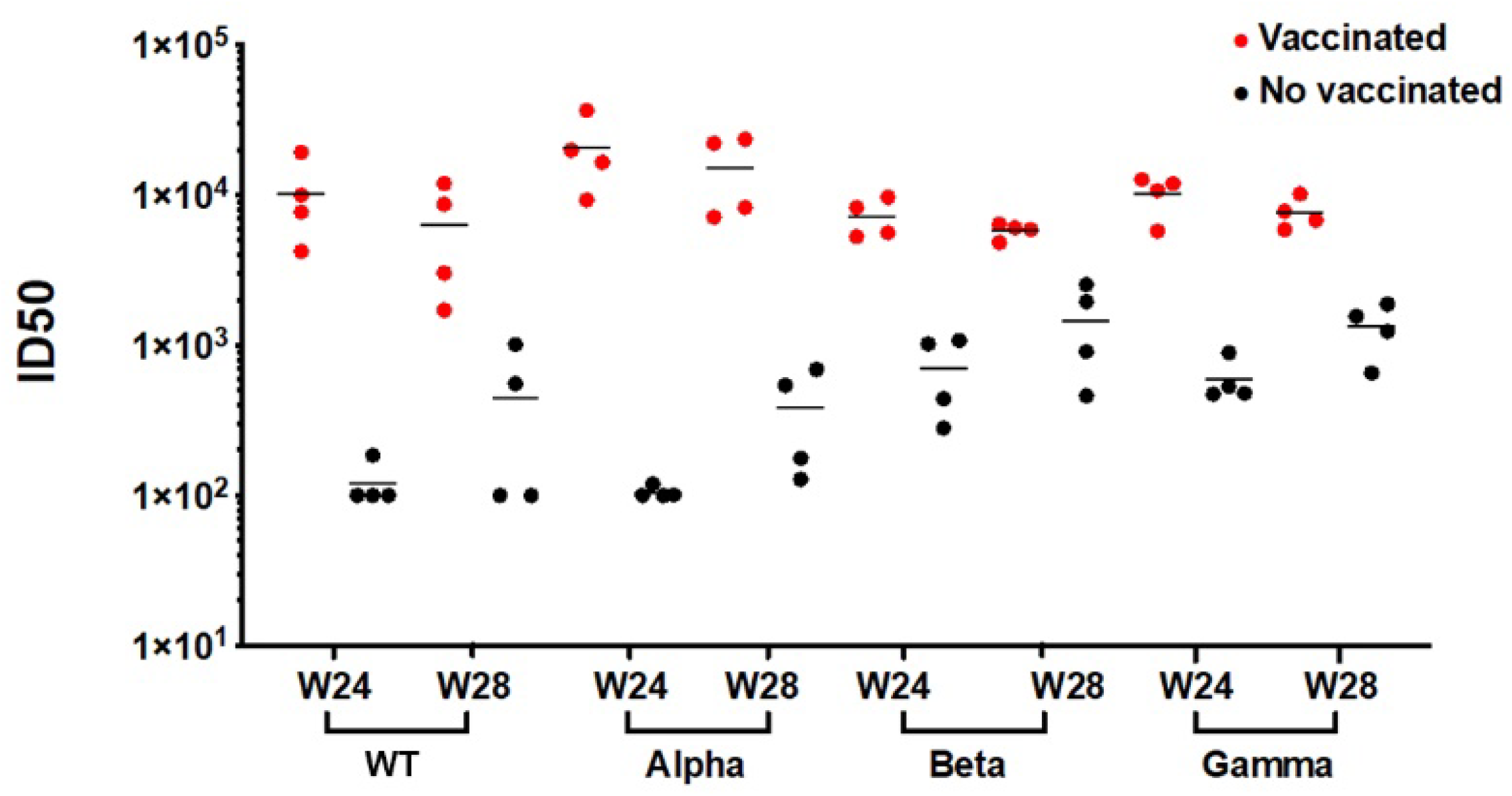
S-VLP vaccination induces neutralization of SARS CoV-2 variants. B.1.1.7 (Alpha, UK), B.1.351 (Beta, SA) and P.1 (Gamma, BR) pseudovirus neutralization titers were compared to the Wuhan vaccine strain (WT). Titers were determined at weeks 24 (exposure) and 28 (4 weeks pe). Comparison of sera from vaccinated macaques and the control group indicated high background values at week 24 (challenge) for Beta and Gamma.

**Figure S5:**
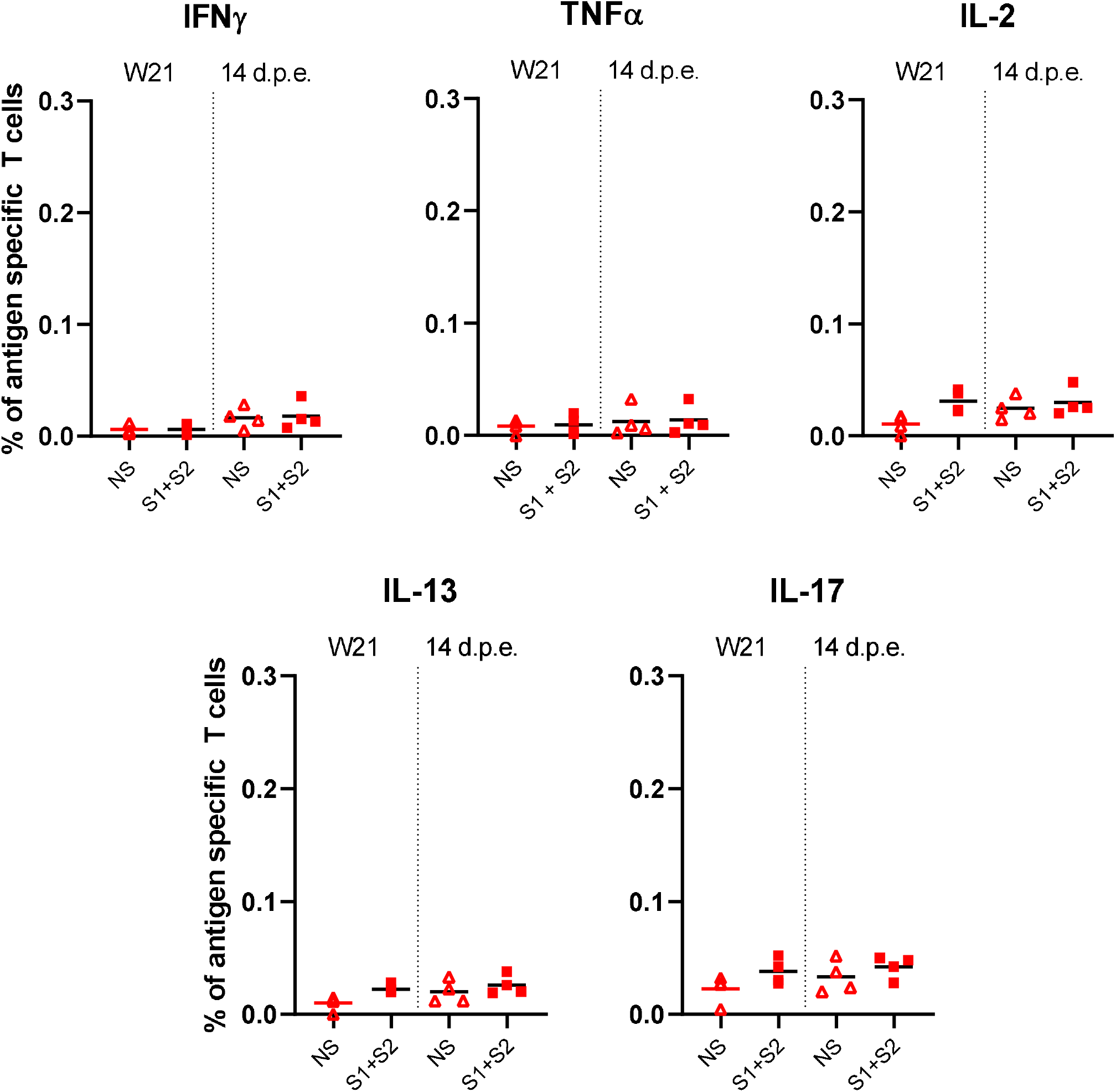
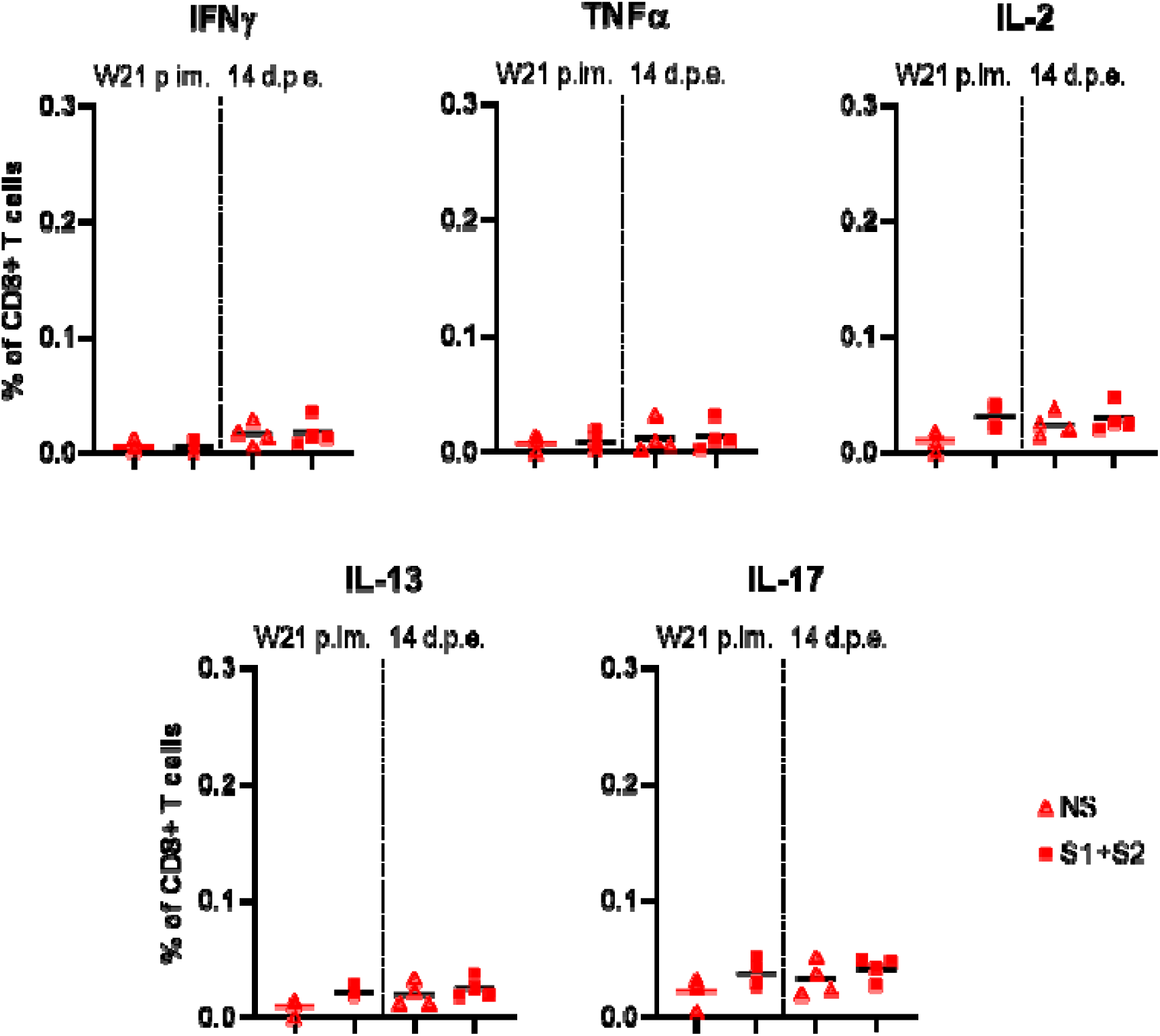
Antigen-specific CD8 T-cell responses in S-VLP immunized cynomolgus macaques. Frequency of IFNγ+ (top left), TFNα+ (top middle), IL-2+ (top right), IL-13+ (bottom left) and IL-17+ (bottom right) antigen-specific CD8+ T cells (CD137+) in the total CD8+ T cell population, respectively, for each immunized macaque (n = 4) at week (W)21 (i.e. two weeks after the 4^th^ immunization, pre-exposure) and 14 days post-exposure (d.p.e.). PBMCs were stimulated overnight with medium (open symbols) or SARS-CoV-2 S overlapping peptide pools (filled symbols). Time points in each experimental group were compared using the Wilcoxon signed rank test.

**Figure S6:**
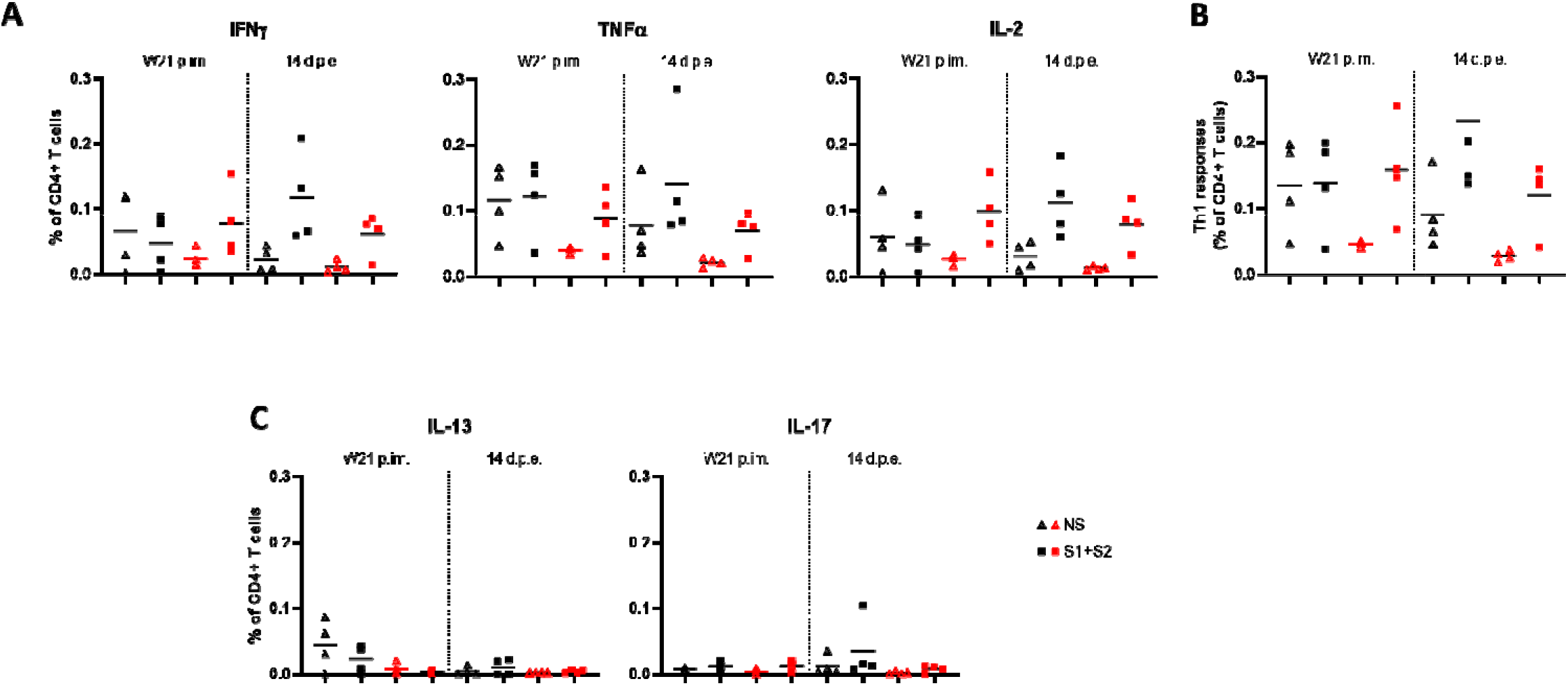
Antigen-specific CD4 T-cell responses in cynomolgus macaques. Frequency of (A) IFNγ+, TFNα+ and IL-2+, (B), Th1 (IFN γ+/−, IL-2+/−, TNFα+), (C) IL-13+and IL-17+ antigen-specific CD4+ T cells (CD154+) in the total CD4+ T cell population, respectively, for each control (n=4, black) and immunized macaque (n = 4, red) at week (W)21 post-first immunization (p.im.) (i.e. two weeks after the 4^th^ immunization, pre-exposure) and 14 days post-exposure (d.p.e.). PBMCs were stimulated overnight with medium (open symbols) or SARS-CoV-2 S overlapping peptide pools (filled symbols). Time points in each experimental group were compared using the Wilcoxon signed rank test. Groups were compared using the non-parametric Mann-Whitney test.

